# Acylation of the incretin peptide exendin-4 directly impacts GLP-1 receptor signalling and trafficking

**DOI:** 10.1101/2021.04.01.438030

**Authors:** Maria Lucey, Tanyel Ashik, Amaara Marzook, Yifan Wang, Joëlle Goulding, Atsuro Oishi, Johannes Broichhagen, David J Hodson, James Minnion, Yuval Elani, Ralf Jockers, Stephen J Briddon, Stephen R Bloom, Alejandra Tomas, Ben Jones

## Abstract

The glucagon-like peptide-1 receptor (GLP-1R) is a class B G protein-coupled receptor and mainstay therapeutic target for the treatment of type 2 diabetes and obesity. Recent reports have highlighted how biased agonism at the GLP-1R affects sustained glucose-stimulated insulin secretion through avoidance of desensitisation and downregulation. A number of GLP-1R agonists (GLP-1RAs) feature a fatty acid moiety to promote albumin binding in order to prolong their pharmacokinetics, but the potential for these ligand changes to influence GLP-1R signalling has rarely been investigated beyond potency assessments for cyclic adenosine monophosphate (cAMP). In this work we directly compare the prototypical GLP-1RA exendin-4 with its C-terminally acylated analogue, exendin-4-C16, for their relative propensities to recruit and activate G proteins and β-arrestins, endocytic and post-endocytic trafficking profiles, and interactions with model and cellular membranes. Both ligands had similar cAMP potency but the exendin-4-C16 showed ∼2.5-fold bias towards G protein recruitment and a ∼60% reduction in β-arrestin-2 recruitment efficacy compared to exendin-4, as well as reduced GLP-1R endocytosis and preferential targeting towards recycling pathways. These effects were associated with a reduced ability to promote the movement of the GLP-1R extracellular domain, as determined using a conformational biosensor approach, and a ∼70% increase in insulin secretion. Interactions with plasma membrane lipids were enhanced by the acyl chain. Exendin-4-C16 showed extensive albumin binding and was highly effective for lowering of blood glucose in mice over at least 72 hours. Overall, our study highlights the importance of a broad approach to the evaluation of GLP-1RA pharmacology.

**Significance statement:** Acylation is a common strategy to enhance the pharmacokinetics of peptide-based drugs. Our work shows how acylation can also affect various other pharmacological parameters, including biased agonism, receptor trafficking and interactions with the plasma membrane, which may be therapeutically important.

## 1 Introduction

Type 2 diabetes (T2D) affects over 400 million people worldwide and leads to chronic illnesses including blindness, nerve damage, kidney failure and cardiac disease (Guariguata *et al*., 2014). Pathological increases in blood glucose, the central metabolic defect in T2D, occur chiefly due to failure of pancreatic beta cells to release enough of the glucoregulatory hormone insulin, and overweight-related tissue resistance to insulin action. Therefore, pharmacological targeting of the glucagon-like peptide-1 receptor (GLP-1R), a class B G protein-coupled receptor (GPCR) which increases insulin release in a glucose-dependent manner and also suppresses appetite leading to weight loss, is an effective strategy against T2D (Graaf *et al*., 2016). As GLP-1(7-36)NH_2_, the endogenous ligand for GLP-1R, is enzymatically inactivated in the circulation within minutes and eliminated by glomerular filtration, a number of strategies have been used to develop pharmaceutically viable peptide GLP-1R agonists (GLP-1RAs) which remain active in the circulation for hours or days (A Andersen *et al*., 2018). As typified by exendin-4 (in clinical use as exenatide), the peptide amino acid sequence can be altered to prevent proteolytic degradation, extending circulatory half-life from minutes to hours, whilst retaining full activity at the receptor (Kolterman *et al*., 2003). An alternative or complementary approach, the best known exemplars of which are liraglutide and semaglutide (Lau *et al*., 2015), involves attachment of a fatty acid side chain to the peptide to allow it to bind reversibly to albumin. As albumin exceeds the size limit for glomerular filtration, this major elimination route is lost (Malm-Erjefält *et al*., 2010). Half-life protraction of semaglutide via this strategy allows once-weekly administration in humans.

Whilst GLP-1RA pharmacology has been traditionally assessed by measuring cyclic adenosine monophosphate (cAMP) responses, additional factors are now recognised as important determinants of insulin release. For example, changes to the amino acid sequence of GLP-1R peptide agonists in their receptor core-facing N-terminal region can lead to bias between G protein signalling, β-arrestin recruitment and receptor endocytosis (H Zhang *et al*., 2015; Jones *et al*., 2018). We recently reported how a dual strategy based on both pharmacokinetically preferential acylation and G protein-directed biased agonism could optimise GLP-1R therapeutic efficacy (Lucey *et al*., 2020). Whilst not explored in detail in the latter project, we observed that the addition of a C-terminal acyl chain further influenced the ability of each ligand to recruit G proteins and β-arrestins to the GLP-1R, highlighting this feature as a modifier of GLP-1R efficacy in its own right.

Moreover, it is conceivable that acylation could constructively influence the interaction between agonist and plasma membrane by conferring distinct physicochemical properties, as suggested for the acylated analogues of glucagon-like peptide-2 (GLP-2) (Trier *et al*., 2014). This has the potential to modulate downstream signalling by directing the ligand towards GLP-1Rs situated in sub-microscopic membrane domains variably enriched with signalling effectors on the cytoplasmic side (Villar *et al*., 2016). Indeed, differences in segregation of activated GLP-1Rs into these “membrane rafts” are an important feature of biased agonist action (Buenaventura *et al*., 2019).

To investigate these possibilities, in this study we performed a detailed comparison of the prototypical GLP-1R therapeutic agonist exendin-4 with an equivalent peptide bearing a C16 fatty diacid at its C-terminus, focusing on the relative propensities of each ligand to engage with G protein and β-arrestin recruitment and activation, endocytosis and post-endocytic sorting. We also evaluated interactions made by each ligand with giant unilamellar vesicles (GUVs) and the relative impact of modulating cellular cholesterol levels on GLP-1R signalling with each ligand. Our study highlights how installation of an acyl chain at the C-terminus influences multiple pharmacological properties of exendin-4, enhancing its ability to stimulate insulin release from pancreatic beta cells.

## 2 Materials and methods

### 2.1 Peptides and reagents

All peptides were obtained Wuxi Apptec (Wuhan, China) and were at least 90% pure. Laboratory reagents were obtained from Sigma Aldrich (Dorset, UK) unless otherwise specified.

### 2.2 Cell culture

HEK293T cells were maintained in DMEM supplemented with 10% FBS and 1% penicillin/streptomycin. HEK293-SNAP-GLP-1R cells generated by stable transfection of pSNAP-GLP-1R (Cisbio, Paris, France) into HEK293 cells (Buenaventura *et al*., 2019) were used for some experiments, and maintained in DMEM supplemented with 10% FBS, 1% penicillin/streptomycin and 1 mg/ml G418. T-REx-SNAP-GLP-1R cells (Fang, Chen, Manchanda, *et al*., 2020), generated from parental Flp-In™ T-REx™ 293 cells (Thermo Fisher Scientific, MA, USA) by Flp-mediated genomic integration of an insert encoding human GLP-1R with an N-terminal SNAP_f_-tag and C-terminal SmBiT tag (custom lingen, Germany), with pcDNA5/FRT as the backbone, were maintained in DMEM supplemented with 10% FBS and 1% penicillin/streptomycin. SNAP-GLP-1R expression was induced using 0.1 µg/ml tetracycline for 24 hours prior to experiments. INS-1 832/3 cells (a gift from Prof Christopher Newgard, Duke University) (Hohmeier *et al*., 2000) were maintained in RPMI at 11 mM glucose, supplemented with 10% FBS, 10 mM HEPES, 1 mM pyruvate, 1 mM pyruvate, 50 μM β-mercaptoethanol and 1% penicillin/streptomycin.

### 2.3 Equilibrium GLP-1R binding assays

HEK293-SNAP-GLP-1R cells were labelled using 40 nM SNAP-Lumi4-Tb (Cisbio) in complete medium for 60 min at room temperature. After washing, cells were resuspended in HBSS containing 0.1% BSA and metabolic inhibitors (10 mM NaN_3_ and 20 mM 2-deoxyglucose) to prevent endocytosis (Widmann *et al*., 1995). After 20 min at room temperature, cells in 96-well white plates were equilibrated to 4°C before addition of agonists. To measure binding parameters for exendin-4 and exendin-4-C16, these non-labelled ligands were added alongside 10 nM exendin(9-39)-FITC and incubated for 24 hours in the dark to allow binding equilibrium to be achieved. TR-FRET signal from each well was then recorded using a Flexstation 3 with the following settings: λ_ex_ 340 nm, λ_em_ 520 and 620 nm, auto-cutoff, delay 50 µs, integration time 300 µs. A saturation binding isotherm for exendin(9-39)-FITC was performed during each experiment, and the K_i_ values for competing agonists were determined using the “Fit K_i_” preset of Prism 8.0. To measure saturation binding for FITC- or TMR-labelled exendin-4 and exendn-4-C16 directly, these ligands were added to cells and incubated as above, followed by measurement of TR-FRET signal using the following settings: FITC ligands: λ_ex_ 340 nm, λ_em_ 520 and 620 nm, auto-cutoff, delay 50 µs, integration time 300 µs; TMR ligands: λ_ex_ 340 nm, λ_em_ 550 and 610 nm, auto-cutoff, delay 50 µs, integration time 300 µs.

### 2.4 Measurement of cAMP production

HEK293-SNAP-GLP-1R cells were resuspended in serum-free medium and treated at 37°C with indicated concentration of agonist for 30 min. cAMP was then assayed by HTRF (Cisbio cAMP Dynamic 2) using a Spectramax i3x plate reader (Molecular Devices). Where indicated, cells were pre-incubated with methyl- β-cyclodextrin in advance of the assay before washing.

### 2.5 NanoBiT assay

The assay was performed as previously described (Lucey *et al*., 2020). HEK293T cells in 12-well plates were transfected with 0.5 µg GLP-1R-SmBiT plus 0.5 µg LgBiT-mini-G_s_ (Wan *et al*., 2018) (a gift from Prof Nevin Lambert, Medical College of Georgia, USA), or with 0.05 µg GLP-1R-SmBiT and 0.05 µg LgBit-β-arrestin-2 (Promega, Hampshire, UK) plus 0.9 µg pcDNA3.1 for 24 hours. Cells were detached with EDTA, resuspended in HBSS, and furimazine (Promega) was added at a 1:50 dilution from the manufacturer’s pre-prepared stock. After dispensing into 96-well white plates, a baseline read of luminescent signal was serially recorded over 5 min using a Flexstation 3 instrument (Molecular Devices) at 37°C before addition of the indicated concentration of ligand, after which signal was repeatedly recorded for 30 min. Results were expressed relative to individual well baseline and AUC was calculated. Each assay was performed with both pathways measured in parallel using the same ligand stocks. Bias between mini-G_s_ and β-arrestin-2 recruitment was determined similarly to earlier work (Pickford *et al*., 2020), by subtracting the log /K values (Kenakin *et al*., 2012) for each ligand in each pathway for each assay to determine Δ log /K_A_ (referred to here as Δ LogR); this indicates the relative pathway preference for each ligand which can then be compared statistically.

### 2.6 β-arrestin activation assay

The assay was adapted from a previous description (Oishi *et al*., 2019). HEK293T cells were seeded in 6-well plates and transfected with 0.5 µg NLuc-β-arrestin-2-CyOFP under a TK promoter, 0.5 µg SNAP-GLP-1R under a CMV promoter, plus 1 µg pcDNA3.1, for 24 hours before the assay. Alternatively, stable HEK293-SNAP-GLP-1R cells in 6-well plates were transfected with 0.5 µg NLuc-β-arrestin-2-CyOFP plus 1.5 µg pcDNA3.1. Cells were detached with EDTA, resuspended in HBSS with furimazine (1:50 dilution). After dispensing into 96-well white plates, a baseline read of luminescent signals at both 460 nm and 575 nm was serially recorded over 5 min using a Flexstation 3 instrument at 37°C. Ligands in HBSS were then added, after which signal was repeatedly recorded for 30 min. Results were first normalised to individual well baseline and then to the average vehicle signal. Statistical comparisons were performed on AUC calculated from each ligand-induced kinetic trace.

### 2.7 G protein activation assay by Nb37 BRET

A construct encoding SNAP-GLP-1R with a C-terminal nanoluciferase was generated in house by PCR cloning of the nanoLuciferase sequence from pcDNA3.1-ccdB-Nanoluc (a gift from Mikko Taipale; Addgene plasmid # 87067) onto the C-terminus end of the SNAP-GLP-1R vector (Cisbio), followed by site-directed mutagenesis of the GLP-1R stop codon. HEK293T cells in 6-well plates were transfected with 0.2 µg SNAP-GLP-1R-NLuc, 0.2 µg Nb37-GFP (a gift from Dr Roshanak Irranejad, UCSF), plus 1.6 µg pcDNA3.1, for 24 hours before the assay. Cells were detached with EDTA, resuspended in HBSS, and furimazine was added at a 1:50 dilution from the manufacturer’s pre-prepared stock. After dispensing into 96-well white plates, a baseline read of luminescent signals at both 460 nm and 525 nm was serially recorded over 5 min using a Flexstation 3 instrument at 37°C. Ligands in HBSS were then added, after which signal was repeatedly recorded for 30 min. Results were first normalised to individual well baseline and then to the average vehicle signal. Statistical comparisons were performed on AUC calculated from each ligand-induced kinetic trace.

### 2.8 Preparation and imaging of fixed cell samples

Cells were seeded onto coverslips coated with 0.1% poly-D-lysine and allowed to adhere overnight. Where relevant, transfection was performed as for functional assays and surface labelling of SNAP-tagged GLP-1R was performed using 0.5 µM of the indicated SNAP-Surface probe for 30 min at 37°C before washing with HBSS. Ligands were applied in Ham’s F12 media containing 0.1% BSA at 37°C. For fixation, 4% paraformaldehyde (PFA) was applied directly to the medium for 15 min before washing with PBS. Slides were mounted in Prolong Diamond antifade with DAPI (Thermo Fisher Scientific, MA, USA) and allowed to set overnight. Imaging was performed using a Nikon Ti2E custom microscope platform with automated stage (ASI) and LED light source controlled by µManager. High resolution z-stacks were acquired via a 100X 1.45 NA oil immersion objective, and used for image deconvolution using DeconvolutionLab2 (Sage *et al*., 2017) with the Richardson-Lucy algorithm.

### 2.9 Measurement of GLP-1R internalisation by time-lapse high content microscopy

HEK293-SNAP-GLP-1R cells were seeded overnight in black, clear-bottom 96-well imaging plates coated with 0.1% poly-D-lysine. Cells were labelled with SNAP-Surface-649 (0.5 µM, from New England Biolabs, Hitchin, UK) for 30 min at 37°C. After washing, imaging medium was added to the wells (phenol red-free DMEM with 10 mM HEPES and 0.1% BSA) at 37°C. Baseline epifluorescence and transmitted phase contrast images were acquired using a 20X magnification 0.75 NA objective from 4 positions per well using the imaging system in Section 2.8. Further baseline epifluorescence images were acquired using a 40X magnification 0.95 NA objective, after which ligands were added directly to each well. Images in the same 4 positions were acquired every 3 min for 30 min. To quantify the translocation of surface-labelled SNAP-GLP-1R from plasma membrane to endosomes, a “spot-counting” imaging processing pipeline implemented in Fiji was developed as follows: 1) flatfield correction was applied to the epifluorescence images using BaSiC (Peng *et al*., 2017); 2) Laplacian filtering (smoothening scale: 2.0) was applied using FeatureJ; 3) starting with an image in which multiple endosomal puncta were visible, images were thresholded using the auto-selected threshold from the “Triangle” algorithm, and the same threshold was used for all images; 4) spots were counted using the particle counting algorithm in Fiji with a size limit of 0.5-5 µm, and roundness of 0.5-1; 5) differences in cell density within each image were accounted for by estimating confluence from the phase contrast image (cropped to the region covered by the 40X objective) using PHANTAST (Jaccard *et al*., 2014) and dividing the spot count by this value; 6) after normalisation, the number of spots present at baseline was subtracted to identify the ligand-induced changes. Rate constants were determined from one-phase exponential curve fitting in Prism 8.0.

### 2.10 Measurement of GLP-1R internalisation by DERET

The assay was performed as previously described (Lucey *et al*., 2020). HEK-SNAP-GLP-1R cells were labelled using 40 nM SNAP-Lumi4-Tb in complete medium for 60 min at room temperature. After washing, cells were resuspended in HBSS containing 24 µM fluorescein and dispensed into 96-well white plates. A baseline read was serially recorded over 5 min using a Flexstation 3 instrument at 37°C in TR-FRET mode with the following settings: λ_ex_ 340 nm, λ_em_ 520 and 620 nm, auto-cutoff, delay 400 µs, integration time 1500 µs. Ligands were then added, after which signal was repeatedly recorded for 30 min. Fluorescence signals were expressed ratiometrically after first subtracting signal from wells containing 24 µM fluorescein without cells. Internalisation was quantified as AUC relative to individual well baseline.

### 2.11 Measurement of receptor recycling by TR-FRET

The assay was performed as previously described (Fang, Chen, Pickford, *et al*., 2020). Adherent HEK293-SNAP-GLP-1R cells in 96-well white plates coated with 0.1% poly-D-lysine were labelled with 40 nM BG-S-S-Lumi4-Tb in complete medium for 60 min at room temperature. After washing, cells were treated for 30 min with each agonist at 100 nM or vehicle (serum-free medium) to induce GLP-1R internalisation. Cells were then washed once with HBSS before application of alkaline TNE buffer (pH 8.6) containing (or not) 100 mM Mesna for 5 min at 4°C to cleave residual surface GLP-1R. 10 nM Luxendin645 (Ast *et al*., 2020) in HBSS containing 0.1% BSA was then added and TR-FRET signal was serially monitored using a Spectramax i3x multi-mode plate reader with HTRF module. Ratiometric HTRF responses were normalised to baseline from the first 3 reads.

### 2.12 Measurement of GLP-1R compartmentalization by BRET

HEK293T cells in 6-well plates were transfected with 0.2 µg SNAP-GLP-1R-NLuc, 0.2 µg of KRAS-Venus, Rab5-Venus, Rab7-Venus or Rab11-Venus (kindly provided by Prof Nevin Lambert, Medical College of Georgia, and Prof Kevin Pfleger, University of Western Australia), plus 1.6 µg pcDNA3.1, 24 hours before the assay. Cells were detached with EDTA, resuspended in HBSS, and furimazine was added at a 1:50 dilution from the manufacturer’s pre-prepared stock. After dispensing into 96-well white plates, a baseline read of luminescent signals at both 460 nm and 535 nm was serially recorded over 5 min using a Flexstation 3 instrument at 37°C. Ligands in HBSS were then added, after which signal was repeatedly recorded for 30 min. Results were first normalised to individual well baseline and then to the average vehicle signal. Statistical comparisons were performed on AUC calculated from each ligand-induced kinetic trace.

### 2.13 Mini-G_s_ subcellular translocation assay

HEK293-SNAP-GLP-1R cells were seeded in 6-well plates and transfected with 0.2 µg plasmid DNA encoding mini-G_s_-NLuc (a gift from Prof Nevin Lambert, Medical College of Georgia) (Wan *et al*., 2018), 0.2 µg KRAS-Venus or Rab5-Venus, plus 1.6 µg pcDNA3.1, for 24 hours before the assay. Cells were detached with EDTA and resuspended in HBSS with furimazine (1:50 dilution). After dispensing into 96-well white plates, a baseline read of luminescent signals at both 460 nm and 535 nm was serially recorded over 5 min using a Flexstation 3 instrument at 37°C. Ligands in HBSS were then added, after which signal was repeatedly recorded for 30 min. Results were first normalised to individual well baseline and then to the average vehicle signal. Statistical comparisons were performed on AUC calculated from each ligand-induced kinetic trace.

### 2.14 Insulin secretion assay

INS-1 832/3 cells were pre-incubated in complete medium at 3 mM glucose for 16 hours prior to the assay. After washing, cells were dispensed into 96-well plates containing 11 mM glucose ± the indicated concentration of agonist and incubated for 16-18 hours. Secreted insulin was analysed from a sample of diluted supernatant by immunoassay (Cisbio wide range HTRF assay, measured using a Spectramax i3x plate reader). Results were normalised to the glucose-only response.

### 2.15 Isolation and imaging of giant unilamellar vesicles

GUVs in the fluid phase, composed of 1-palmitoyl-2-oleoyl-sn-glycero-3-phosphocholine (POPC) lipid were prepared via electroformation. Lipids were purchased from Avanti Polar Lipids (AL, USA). First, lipid in chloroform (20_μl; 1_mg/mL) was spread evenly on a conductive indium tin oxide coated (ITO) slide, leaving a film which was dried under vacuum for 30_min to remove residual solvent. A 5_mm thick polydimethylsiloxane (PDMS) spacer with a central cut-out was used to separate the slides with the conductive sides facing each other, and the chamber was filled with a solution of 100 mM sucrose in DI water. An alternating electric field (1_V, 10_Hz) was applied across the ITO plates using a function generator (Aim-TTi, TG315). After 2_h, the electric field was changed to 1_V, 2_Hz for a further hour, and the resulting vesicles collected. Either acylated or non-acylated compounds were added to make a final concentration of 100 nmol/l, and the solution incubated for 30 min. For imaging, 10 ul of the vesicle suspension was added to 90 ul of 100 mM glucose in DI water in an imaging chamber, and the vesicles sedimented on the bottom of the glass slide. Confocal imaging was done on a Leica TCS SP5 confocal fluorescent microscope, with a 20X objective set with an 84.5 µm pinhole (1 airy unit). Samples were acquired at a frequency of 400 Hz with eight line averages. The excitation was achieved a wavelength of 495 nm with emission set at between 510 and 530 nm. The images were acquired in the midplane of the GUVs. During data collection, focal planes slightly above and below were viewed to confirm image acquisition from the midplane.

### 2.16 Peptide-membrane interaction assay by FRET

T-REx-SNAP-GLP-1R cells were pre-treated ± 0.1 µg/ml tetracycline to induce SNAP-GLP- 1R expression. Cells were detached using EDTA, centrifuged, and the pellet washed before the assay. Cellular labelling was then performed for 5 min using freshly prepared NR12S (Kucherak *et al*., 2010), a gift from Prof Andrey Klymchenko, University of Strasbourg, at the indicated concentration in PBS in the dark. After washing three times to remove unbound probe, cells were dispensed into black 96-well microplates containing FITC-conjugated peptides at a final concentration of 100 nM. FRET measurements were taken using a Flexstation 3 instrument with the following settings: λ_ex_ 475 nm, λ_em_ range 520 – 680 nm in 5 nm increments, cutoff 495 nm. Recorded fluorescence intensities were then normalised to 525 nm to allow for well-to-well differences and signal quenching by FRET. The normalised spectral trace from wells containing cells without NR12S probe was subtracted to determine probe-related FRET signal, which was quantified as AUC between 525 and 680 nm. Gaussian fitting was performed to determine the peak emission wavelength and relative intensity.

### 2.17 GLP-1R conformational sensor assay

HEK293-SNAP-GLP-1R cells were labelled with Lumi4-Tb (40 nM, 60 min at 37°C, in complete medium). After washing, cells were resuspended in HBSS containing NR12A (50 nM) and seeded into half-area opaque white plates. Baseline signal was measured for 5 min at 37°C using a Flexstation 3 plate reader with the following settings: λ_ex_ = 335 nm, λ_em_ = 490 and 590 nm, delay 50 μ s. Ligands were then added, and signal was serially monitored for 10 min. The TR-FRET ratio, i.e. the ratio of fluorescence intensities at 590 and 490 nm, was considered indicative of the proximity of the GLP-1R extracellular domain (ECD) to the plasma membrane.

### 2.18 Fluorescence correlation spectroscopy (FCS)

FCS was performed as previously described (Corriden *et al*., 2014) on a Zeiss LSM 510NLO ConfoCor 3 microscope fitted with a 40X c-Apochromat 1.2 NA water-immersion objective. 30 s reads were taken at 22°C with a Diode-pumped solid-state (DPSS) 561nm laser excitation at ∼0.2 kWcm^-2^, emission collected through a 580-610nm bandpass and the pinhole set to 1 airy unit. The measurement volume was calibrated on each experimental day with 20 nM Rhodamine 6 G (Diffusion Coefficient, D, 2.8 x10^-10^ m^2^s^-1^) in HPLC grade water. Exendin-4-TMR and Exendin-4-TMR-C16 were prepared in HBSS with or without 0.1% BSA in an 8-well Nunc™ Lab-tek™ chambered coverglass (No. 1.0 borosilicate glass bottom) with the measurement volume placed 200 µm above the coverslip bottom. Autocorrelation curves were modelled in Zen 2012 (Carl Zeiss, Jena) to describe a single 3D diffusing component under free diffusion (Corriden 2014). To determine the average molecular brightness of the labelled peptide, measurement reads were reanalyzed by Photon Counting Histogram (PCH) analysis within Zen 2012 (Huang *et al*., 2004). PCH analysis can quantify the average molecular brightness, photon counts per molecule, of a species and also provides a means to separate and quantify the concentration of two species which differ in their average molecular brightness. The laser beam profile was approximated to follow a 3D Gaussian distribution, measurement reads were binned at 20 µs, appropriate for fluorescent species in solution, and the first order correction was calculated daily following calibration with 20 nM Rhodamine 6 G.

### 2.19 *In vivo* study

Animals were maintained in specific pathogen-free facilities, with *ad lib* access to food (except prior to fasting studies) and water. Studies were regulated by the UK Animals (Scientific Procedures) Act 1986 of the U.K. and approved by Imperial College London (Project License PB7CFFE7A). Male C57Bl/6 mice (Charles River, UK) were fed a 60% high fat diet (D12492, Research Diets, USA) for 3 months prior to the study to induce obesity and glucose intolerance. The study began after an additional 1-week acclimatisation period during which mice were singly housed and received sham intra-peritoneal (IP) injections. Mice were randomly allocated to treatment, with average group weight confirmed to be similar post-randomisation. The study began at the beginning of the dark phase with a single dose of each agonist or vehicle, combined with an IP glucose tolerance test (IPGTT).

Agonist was prepared in 20% glucose solution at a volume to provide the indicated weight-adjusted agonist dose and 2 g/kg glucose. Blood glucose was monitored before and at 20-min intervals after agonist/vehicle/glucose administration using a hand-held glucose meter. After 72 hours, the IPGTT was repeated at the beginning of the dark phase without further agonist administration. Food intake was assessed during the study by weighing of food at set intervals. Changes to body weight were also assessed at set intervals.

### 2.20 Experimental design and statistical analysis considerations

A preliminary finding that exendin-4-C16 shows reduced β-arrestin-2 recruitment compared to exendin-4 (Lucey *et al*., 2020), gave us a reasonable expectation that the two ligands would display alterations to other aspects of their pharmacology. Therefore we adopted an approach in which we planned to perform at least 5 independent replicates for quantitative cell culture assays as recommended by some authorities (Curtis *et al*., 2018). We did not perform a formal power calculation. In some instances, we observed more subtle differences between compounds that required additional repeats to gain confidence in the magnitude of the effects size, which were then performed to allow comparisons between assays evaluating similar phenomena (e.g. BRET *versus* TR-FRET assays). Animal experiments were performed with 7-8 mice/group without a formal power calculation, but on the basis of our prior experience in determining meaningful differences between GLP-1RAs. The investigators were not fully blinded for *in vivo* studies but did not have reference to treatment allocation whilst conducting experiments; *in vitro* experiments were unblinded. No mice were excluded from the analyses. In cell culture experiments, technical replicates were averaged so that each individual experiment was treated as one biological replicate. Quantitative data were analysed using Prism 8.0 (GraphPad Software). Dose responses were analysed using 3 or 4-parameter logistic fits, with constraints imposed as appropriate. Bias analyses were performed as described in Section 2.5. Statistical comparisons were made by t-test or ANOVA as appropriate, with paired or matched designs used depending on the experimental design. Mean ± standard error of mean (SEM), with individual replicates in some cases, are displayed throughout. Statistical significance was inferred if p<0.05, without ascribing additional levels of significance.

## 3 Results

### 3.1 Exendin-4-C16 is a G protein-biased ligand at GLP-1R

An exendin-4 analogue with a C16 diacid at the C-terminus with a GK linker, originally described in our earlier publication (Lucey *et al*., 2020), was used in the present study and is referred to as “exendin-4-C16” (Figure 1A). Competition binding experiments with FITC-conjugated antagonist ligand exendin(9-39) in HEK293 cells stably expressing SNAP-GLP-1R (Fang, Chen, Pickford, *et al*., 2020) indicated an approximately two-fold reduction in binding affinity for exendin-4-C16 compared to unmodified exendin-4 (Figure 1B, Table 1). A minor reduction in cAMP signalling potency that was not statistically significant was also observed (Figure 1C, Table 1). To investigate the possibility that exendin-4-C16 may display altered preference for coupling to intracellular effectors, i.e. biased agonism, we used nanoluciferase complementation to measure recruitment of mini-G_s_ (Wan *et al*., 2018) and β arrestin-2 (Dixon *et al*., 2016) to the GLP-1R. This assay indicated that efficacy for β arrestin-2 recruitment was particularly reduced with exendin-4-C16, with bias quantification using a standard approach (van der Westhuizen *et al*., 2014) confirming preferential coupling to mini-G_s_ recruitment (Figure 1D, Table 1).

**Figure 1.**
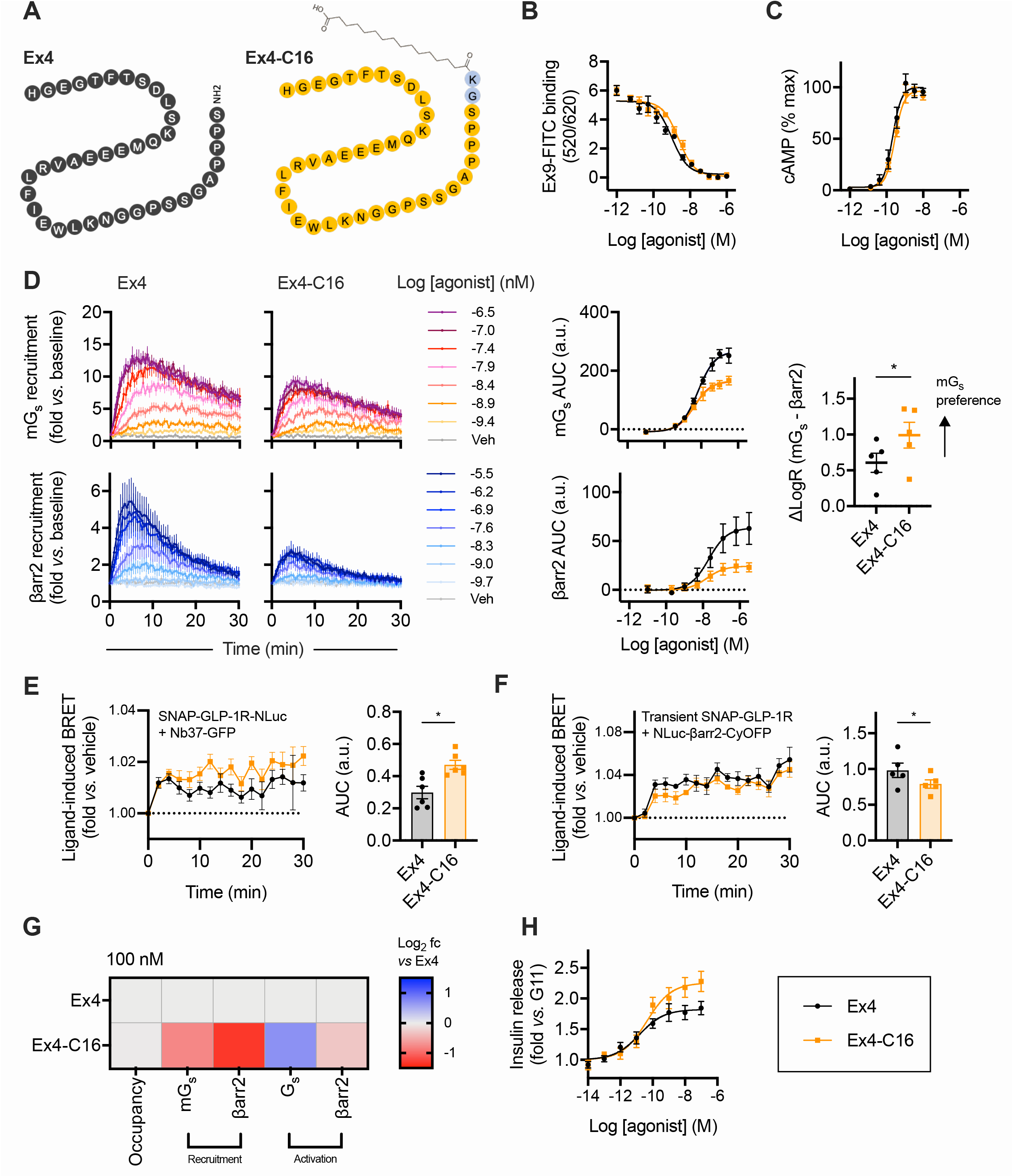
Signal bias with exendin-4 and exendin-4-C16. (**A**) Schematic depicting the amino acid sequences of exendin-4 and exendin-4-C16 in single letter amino acid code. (**B**) Equilibrium binding of unmodified exendin-4 and exendin-4-C16, *n*=5, measured by competition binding in HEK293-SNAP-GLP1-R cells with exendin(9-39)-FITC used as the competing probe. (**C**) cAMP response in HEK293-SNAP-GLP-1R cells, 30 min stimulation, *n*=5, with 4-parameter fits shown after normalisation to global maximum. (**D**) LgBiT-mini-G_s_ (mG_s_) and LgBit-β-arrestin-2 (βarr2) recruitment to GLP-1R-SmBiT, *n*=5, with 3-parameter concentration-responses constructed from AUCs, and bias factor calculation (see methods) with comparison by paired t-test. (**E**) Measurement of G activation in HEK293T cells transiently transfected with SNAP-GLP-1R-NLuc and Nb37-GFP and stimulated with 100 nM agonist or vehicle, *n*=6, with AUCs compared by paired t-test. (**F**) Measurement of β-arrestin-2 activation in HEK293T cells transiently transfected with SNAP-GLP-1R and NLuc-4myc-βarr2-CYOFP1 and stimulated with 100 nM agonist or vehicle, *n*=5, with AUCs compared by paired t-test. (**G**) Heatmap representation of 100 nM agonist response data shown in Figure 1B-F with normalisation to exendin-4 response. (**H**) Insulin secretion in INS-1 832/3 cells stimulated for 16 h with each agonist at 11 mM glucose (G11), *n*=6. Data are shown as mean ± SEM with individual replicates shown in some cases. * p<0.05 by statistical test indicated.

**Table 1.**
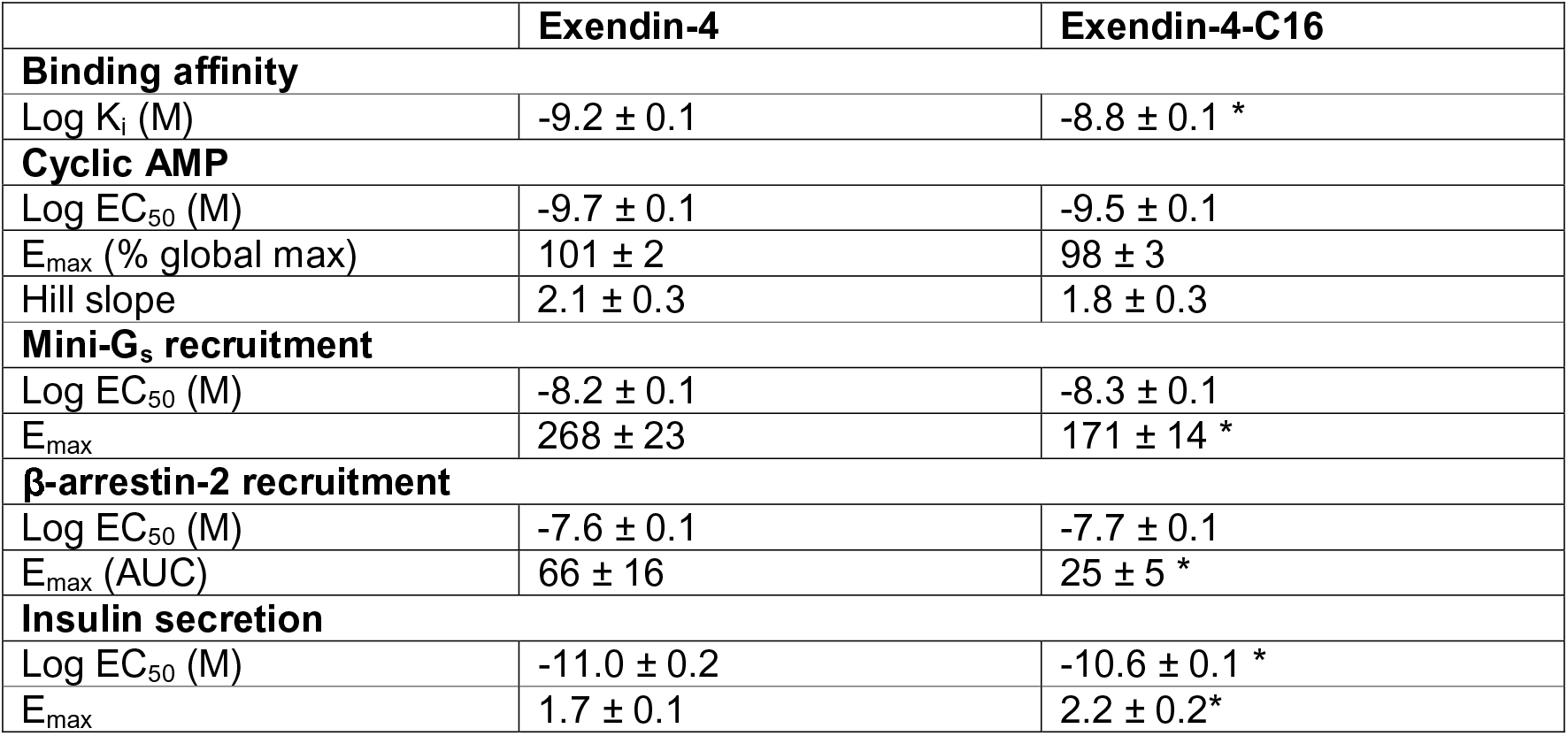
Pharmacological parameters of exendin-4 and exendin-4-C16. Mean ± SEM parameter estimates from concentration-response analyses in Figure 1. * p<0.05 by paired t-test.

|  | **Exendin-4** | **Exendin-4-C16** |
| --- | --- | --- |
| **Binding affinity** |  |  |
| Log $K_i$ (M) | -9.2 ± 0.1 | -8.8 ± 0.1 * |
| **Cyclic AMP** |  |  |
| Log $EC_{50}$ (M) | -9.7 ± 0.1 | -9.5 ± 0.1 |
| $E_{max}$ (% global max) | 101 ± 2 | 98 ± 3 |
| Hill slope | 2.1 ± 0.3 | 1.8 ± 0.3 |
| **Mini-G<sub>s</sub> recruitment** |  |  |
| Log $EC_{50}$ (M) | -8.2 ± 0.1 | -8.3 ± 0.1 |
| $E_{max}$ | 268 ± 23 | 171 ± 14 * |
| **β-arrestin-2 recruitment** |  |  |
| Log $EC_{50}$ (M) | -7.6 ± 0.1 | -7.7 ± 0.1 |
| $E_{max}$ (AUC) | 66 ± 16 | 25 ± 5 * |
| **Insulin secretion** |  |  |
| Log $EC_{50}$ (M) | -11.0 ± 0.2 | -10.6 ± 0.1 * |
| $E_{max}$ | 1.7 ± 0.1 | 2.2 ± 0.2* |

As complementation assays measure effector recruitment but not activation, we also performed further experiments to detect ligand-induced conformational changes in G_s_ and β arrestin-2. For the former we recorded dynamic changes in BRET signal between GLP-1R tagged at the C-terminus with nanoluciferase and GFP-tagged nanobody 37 (Nb37), a genetically encoded intrabody that recognises active G_s_ conformations (Irannejad *et al*., 2013). For the latter we used an intramolecular β-arrestin-2 BRET sensor in which conformational changes lead to an increase in proximity between nanoluciferase at the N-terminus and CyOFP at the C-terminus (Oishi *et al*., 2019). These studies suggsted reduced β-arrestin-2 activation with exendin-4-C16 but, interestingly, increased Gα_s_ activation (Figure 1E, 1F, Supplementary Figure 1A), which could be due to reduced β-arrestin-mediated desensitisation. Ligand pharmacology is summarised in the heatmap shown in Figure 1G, indicating how, at a 100 nM dose to achieve close to 100% receptor occupancy, G_s_ recruitment and activation are favoured compared to β-arrestin-2 responses with exendin-4-C16.

As G protein-directed agonism is now established as a means to improve GLP-1R-mediated insulinotropic efficacy by avoiding receptor desensitisation over prolonged stimulation periods (Jones *et al*., 2018; Fremaux *et al*., 2019; Fang, Chen, Pickford, *et al*., 2020; Lucey *et al*., 2020), we also treated INS-1 832/3 clonal beta cells (Hohmeier *et al*., 2000) for 16 hours with exendin-4 and exendin-4-C16 to measured cumulative insulin secretion. As expected, the maximum response was increased with the -C16 ligand (Figure 1H, Table 1).

### 3.2 Exendin-4-C16 triggers slower GLP-1R internalisation compared to exendin-4

GLP-1R undergoes rapid agonist-induced internalisation (Widmann *et al*., 1995), a process that is intrinsically linked to the spatial orchestration of intracellular signal generation (Fletcher *et al*., 2018; Tomas *et al*., 2019; Manchanda *et al*., 2021). Both exendin-4 and exendin-4-C16 resulted in extensive endocytosis of surface-labelled SNAP-GLP-1R (Figure 2A). To quantify this process, we used high content time-lapse microscopy in which the translocation of surface-labelled SNAP-GLP-1R into the endocytic network is determined from the appearance an intracellular punctate distribution of fluorescent signal (Figure 2B, Supplementary Figure 2A, Supplementary Video 1). Treatment with exendin-4 resulted in faster and more extensive appearance of fluorescent puncta at higher ligand concentrations. Notably, the number of puncta tended to reduce at later time-points with exendin-4 due to the coalescence in a perinuclear location where multiple punctate endosomal structures which could no longer be individually resolved.

**Figure 2.**
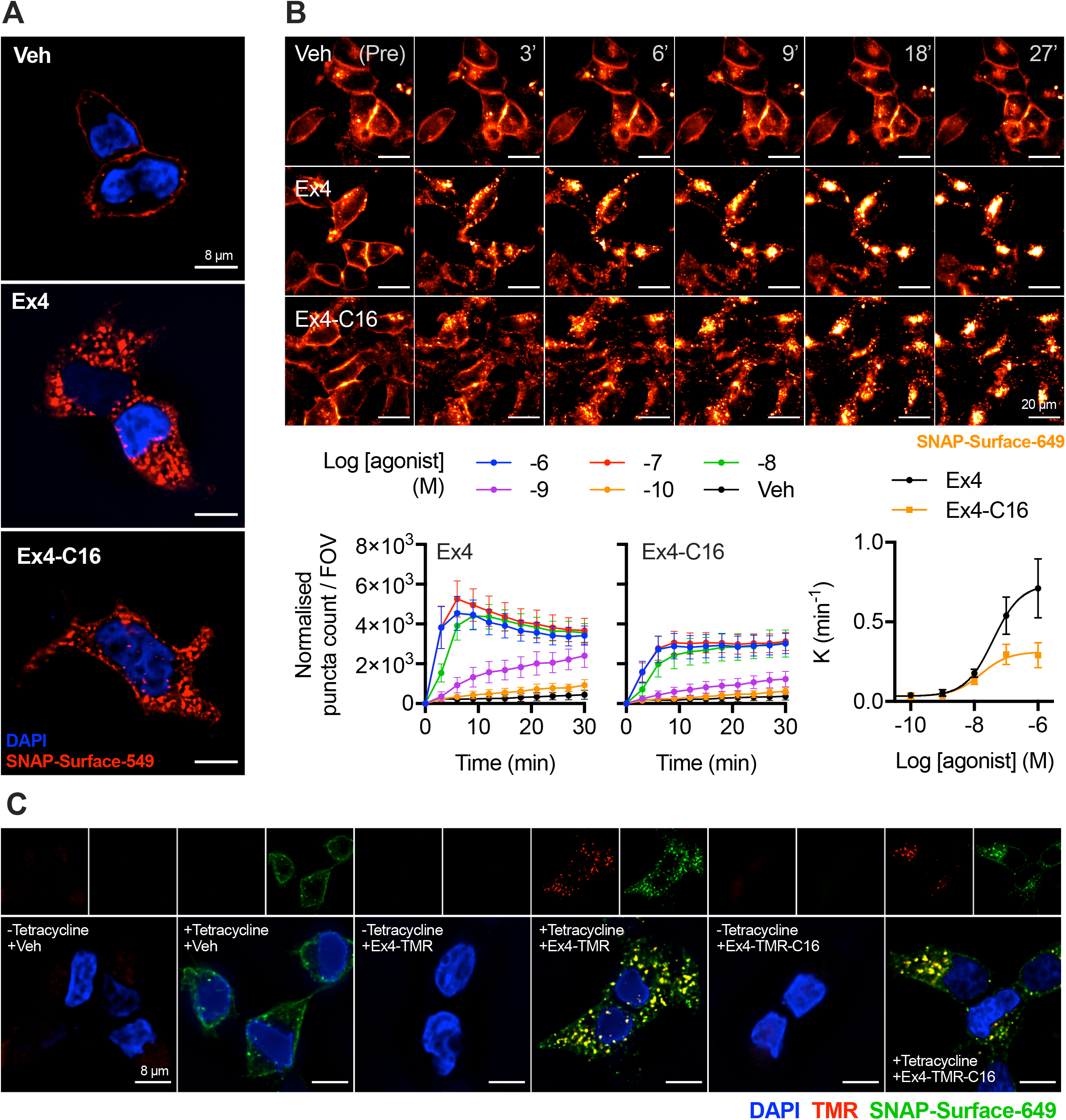
Visualisation of GLP-1R endocytosis with exendin-4 and exendin-4-C16, and fluorescent agonist conjugates. (**A**) Representative images from *n*=3 experiments demonstrating endocytosis of SNAP-GLP-1R labelled with SNAP-Surface-549 after treatment for 30 min with 100 nM agonist or vehicle; scale bars = 8 µm. (**B**) Time-lapse images demonstrating movement of surface-labelled SNAP-GLP-1R into punctate structures on stimulation with 100 nM agonist or vehicle; scale bars = 20 µm. The graphs show the change over time in “spot count” per 0.33 mm field-of-view (FOV) after normalisation to cell confluence, with the concentration-dependent rate constant (K) derived from one-phase association fitting also plotted with a 3-parameter fit. (**C**) Representative images showing endosomal uptake of exendin-4 and exendin-4-TMR in T-REx-SNAP-GLP-1R cells with or without tetracycline-induced GLP-1R expression, labelled with SNAP-Surface-649 prior to stimulation with 100 nM of each TMR-conjugate for 30 min. Scale bars = 8 µm. Data are shown as mean ± SEM.

We also used tetramethylrhodamine (TMR)-tagged conjugates of each ligand, with the fluorophore installed at position K12, previously shown to be well tolerated by exendin-4 (Clardy *et al*., 2014; Jones *et al*., 2018; Pickford *et al*., 2020) and validated for GLP-1R binding by TR-FRET (Supplementary Figure 2B) and cAMP measurements (Supplementary Figure 2C), with the latter showing modest reductions in potency for both TMR-conjugates. Using the tetracycline-inducible T-REx system to modulate SNAP-GLP-1R expression (Fang, Chen, Manchanda, *et al*., 2020), both fluorescent ligands were shown to bind specifically to the receptor and demonstrated extensive uptake into the endosomal compartment (Figure 2C).

### 3.3 Post-endocytic trafficking differences between exendin-4 and exendin-4-C16

To gain insights into ligand-induced GLP-1R movements to or from different subcellular compartments, we used time-resolved proximity-based energy transfer techniques, with both chemical (for FRET) or genetically encoded (for BRET) acceptors as localisation markers, providing complementary readouts. SNAP-GLP-1R labelled at the N-terminus with the lanthanide probe Lumi4-Tb can transfer energy to time-resolved FRET acceptors situated in the extracellular space or the endosomal lumen (Figure 3A). Diffusion-enhanced resonance energy transfer (DERET) was used to quantify movement of GLP-1R away from the cell surface as a reduction in signal transfer to fluorescein-containing extracellular buffer (Levoye *et al*., 2015); this confirmed that GLP-1R internalisation was slower for exendin-4-C16 (Figure 3B, Supplementary Figure 3A). We developed a further assay in which the trafficking of GLP-1R to late endosomes / lysosomes was detected by energy transfer to the lysomotropic fluorescent dye LysoTracker-DND99 (Figure 3C); this suggested that exendin-4-C16 treatment led to markedly less targeting of internalised GLP-1R to this degradative compartment. In contrast, post-endocytic recycling of GLP-1R back to the plasma membrane, measured by time-resolved FRET (Fang, Chen, Pickford, *et al*., 2020) between reemergent GLP-1R and Cy5-labelled antagonist analogue of exendin(9-39) (Ast *et al*., 2020), was faster after treatment with exendin-4-C16 (Figure 3D). TR-FRET data are also shown and statistically compared in Figure 3E.

**Figure 3.**
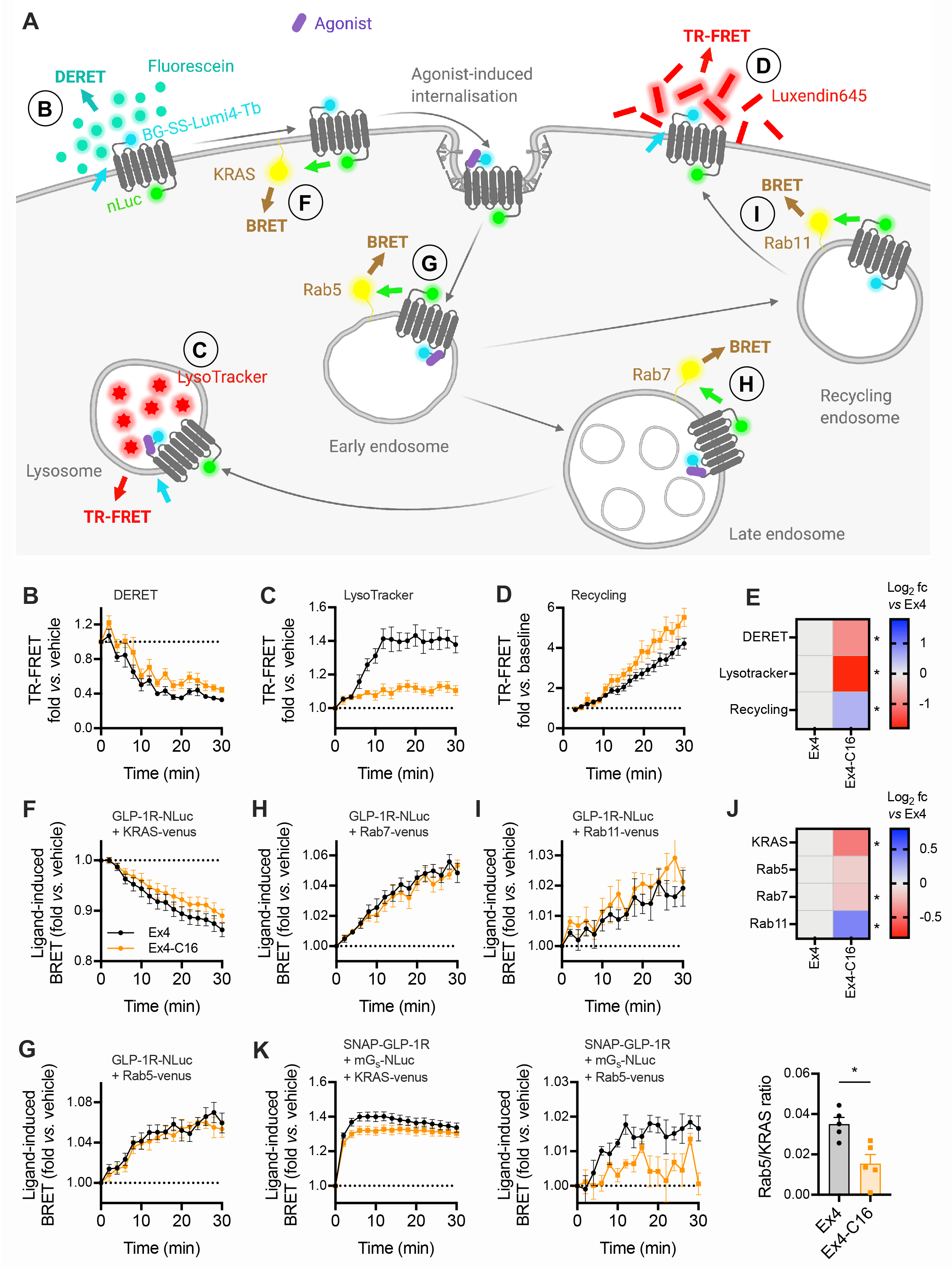
Subcellular targeting of exendin-4 and exendin-4-C16. (**A**) Schematic depicting the investigation of GLP-1R membrane trafficking using the TR-FRET and BRET assays used in this work. (**B**) GLP-1R disappearance from cell surface in HEK293-SNAP-GLP-1R cells treated with 100 nM agonist or vehicle, measured by DERET, *n*=5. (**C**) GLP-1R translocation to late endosomes / lysosomes in HEK293-SNAP-GLP-1R cells treated with 100 nM agonist or vehicle, measured by TR-FRET using Lysotracker Red DND99 as the acceptor, *n*=5. (**D**) GLP-1R recycling in HEK293-SNAP-GLP-1R cells previously treated for 30 min with 100 nM agonist, measured by TR-FRET using extracellular Luxendin645 as the acceptor, *n*=5. (**E**) Heatmap representation of data shown in (B), (C) and (D) after normalisation to the exendin-4 response; statistical significance is indicated after paired t-test. (**F**) GLP-1R disappearance from the cell surface in HEK293T cells transiently transfected with SNAP-GLP-1R-NLuc and KRAS-venus and treated with 100 nM agonist or vehicle, measured by BRET, *n*=6. (**G**) GLP-1R appearance in Rab5+ early endosomes in HEK293T cells transiently transfected with SNAP-GLP-1R-NLuc and KRAS-venus and treated with 100 nM agonist or vehicle, measured by BRET, *n*=6. (**H**) As for (G) but for Rab7+ late endosomes, *n*=7. (**I**) As for (G) but for Rab11+ recycling endosomes, *n*=7. (**J**) Heatmap representation of data shown in (F), (G), (H) and (I) after normalisation to the exendin-4 response; statistical significance is indicated from paired t-tests. (**K**) Mini-G_s_ translocation to the plasma membrane or Rab5+ early endosomes in HEK293-SNAP-GLP-1R cells treated with 100 nM agonist after transient transfection with Mini-G_s_-NLuc plus KRAS-venus or Rab5-venus, *n*=5. The agonist-specific AUC measured for each compartment marker are ratiometrically compared by paired t-test. Data are shown as mean ± SEM with individual replicates shown in some cases. * p<0.05 by statistical test indicated.

Bystander BRET can be used to monitor the translocation of GPCRs within the endocytic network (Tiulpakov *et al*., 2016). We observed a ligand-induced reduction in BRET signal when SNAP-GLP-1R tagged at the C-terminus with nanoluciferase was co-expressed with the plasma membrane marker KRAS-Venus; the effect was more pronounced with exendin-4 than with exendin-4-C16 (Figure 3F). A concomitant increase in BRET signal to Rab5-Venus was seen, indicating entry of the receptor into early endosomes (Figure 3G). The similar Rab5 BRET signal for each ligand in the face of apparently different GLP-1R internalisation rates might be reconciled by the observation that, after exendin-4-C16 treatment, a subtly reduced signal from Rab7-positive late endosomes was detected (Figure 3H), whereas a greater signal was recorded from Rab11-positive recycling endosomes (Figure 3I). BRET results were broadly in accordance with results from the TR-FRET assays, although differences were generally smaller whilst remaining statistically significant (Figure 3J). Thus, exendin-C16 appears to have increased propensity for directing internalised GLP-1R towards a recycling pathway, resulting in a reduced net rate of surface receptor loss.

GLP-1R cAMP signalling was reported to originate from early endosomes as well as the plasma membrane (Girada *et al*., 2017). With nanoluciferase-tagged mini-G_s_ co-expressed with SNAP-GLP-1R, a rapid increase in mini-G_s_-to-KRAS BRET signal was apparent with both ligands (Figure 3K), but with a slightly higher peak response with exendin-4 consistent with the higher efficacy displayed by this ligand for G protein recruitment (Figure 1D). A decline in signal at later time-points was observed with exendin-4 but not exendin-4-C16, which may result from more extensive β-arrestin-mediated steric hinderance or internalisation of GLP-1R with the former ligand. Interestingly, mini-G_s_-to-Rab5 BRET signal amplitude was considerably reduced for exendin-4-C16, despite the fact that the receptor localisation in the early endosomal (EE) compartment was similar for both ligands.

These data indicate that C-terminal acylation alters the trafficking profile of exendin-4, favouring GLP-1R sorting towards a recycling rather than degradative pathway. The observation that mini-G_s_ protein recruitment to EEs was reduced with exendin-4-C16, yet this ligand shows higher efficacy for sustained insulin secretion, suggests that signalling from early endosomes may not be a dominant mechanism for prolongued signalling with GLP-1R under pharmacological conditions, at least with exendin-4 derived GLP-1R agonists.

### 3.4 Acylation affects the interaction of the exendin-4 peptide with the plasma membrane

Several studies report an interaction of GLP-1R agonist peptides with model membranes, typically resulting in enhanced stability of the helical secondary structure (Thornton and Gorenstein, 1994; Neidigh *et al*., 2001; NH Andersen *et al*., 2002; Fox *et al*., 2009). Acylation of glucagon-like peptide-2 increases its tendency to interact with lipid bilayers (Trier *et al*., 2014). Similarly, the C16 chain of exendin-4-C16 might promote engagement with hydrophobic membrane regions. We performed confocal microscopy of FITC conjugates of exendin-4 and exendin-4-C16 bound to giant unilamellar vesicles (GUVs), prepared via electroformation. Both FITC ligands were conjugated at position K12, as for the equivalent TMR conjugates, and retained GLP-1R binding properties (Supplementary Figure 4A) and cAMP signalling (Supplementary Figure 4B). When the acylated compound was present in the sample, clear fluorescence accumulation was seen on the membrane surface, in contrast to the non-acylated compound where no accumulation was seen (Figure 4A). This demonstrates that the presence of the C16 acyl chain promotes insertion of the molecule into model lipid bilayers.

**Figure 4.**
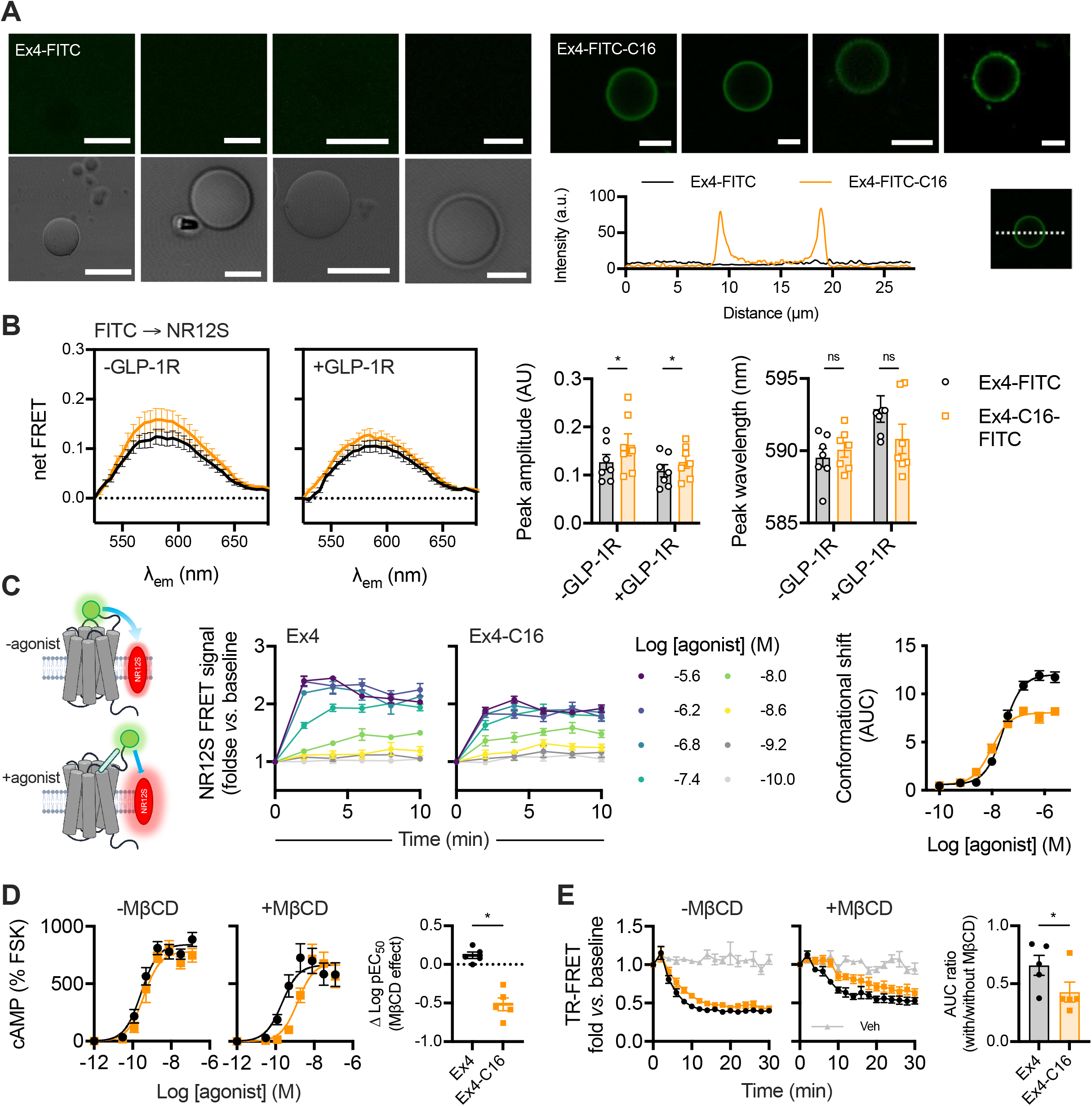
Membrane interactions of C-terminally acylated exendin-4 conjugates. (**A**) Confocal microscopy and phase contrast images of GUVs incubated with exendin-4-FITC or exendin-4-FITC-C16. Scale bars = 5 µm. Membrane signal is represented on the line plot. (**B**) FRET spectrum from T-REx-SNAP-GLP-1R cells with or without tetracycline-induced GLP-1R expression, labelled with 50 nM NR12S and incubated with 100 nM exendin-4-FITC or exendin-4-FITC-C16, *n*=7. The background spectrum of each FITC ligand was subtracted and the trace was normalised to the signal at 525 nm. Peak emission wavelength and amplitude, derived from Gaussian fitting of each spectrum, are shown and compared between ligands by 2-way randomised block ANOVA with Sidak’s test. (**C**) Principle of NR12S conformational biosensor assay, with kinetic responses showing TR-FRET ratio normalised to baseline in response to indicated concentration of exendin-4 or exendin-4- C16, *n*=5, with concentration response curve constructed from AUC with 3-parameter fit. (**D**) cAMP responses in HEK293-SNAP-GLP-1R cells pre-incubated for 30 min with vehicle or 10 mM MβCD, prior to stimulation with each ligand for 30 min. Responses are expressed relative to the forskolin (FSK; 10 µM) response, *n*=5, with the effect of MβCD on potency shown by subtracting pEC_50_ results and comparison by paired t-test. (**E**) GLP-1R internalisation measured by DERET in HEK293-SNAP-GLP-1R cells pre-incubated for 30 min with vehicle or 10 mM MβCD prior to stimulation with each ligand at 100 nM for 30 min, *n*=5. AUC relative to baseline was quantified and +MβCD results expressed relative to -MβCD results for each ligand, and compared by paired t-test. Data are shown as mean ± SEM with individual replicates where possible. * p<0.05 by statistical test indicated.

We also investigated whether the C16-acylated peptide displays increased membrane interactions in living cells by measuring FRET between FITC-ligands and the plasma membrane labelled with NR12S (Kucherak *et al*., 2010). The excitation spectrum of NR12S is well matched to the emission spectrum of FITC and, as a solvatochromic probe, its emission spectrum is polarity-sensitive, potentially allowing discrimination of interactions occurring in membrane domains with different degrees of liquid order, e.g. lipid rafts *versus* non-raft regions. The NR12S spectra obtained from excitation of FITC ligands incubated with NR12S-labelled cells showed an increase in maximum FRET signal without any significant spectral shift (Figure 4B). Similar findings were observed with or without GLP-1R expression in the same cell system. This suggests that exendin-4-C16 may indeed interact with cell membranes to a greater extent than exendin-4, but does not provide evidence that this directs the ligand to interact with specific GLP-1R subpopulations situated in membrane nanodomains with different degrees of liquid order.

To gain further insights into how exendin-4 and exendin-4-C16 interact with GLP-1R at the plasma membrane we used NR12S as a FRET acceptor for the N-terminally SNAP-tagged GLP-1R labelled with Lumi-4-Tb, monitoring dynamic changes in TR-FRET signal with each ligand over time indicative of movement of the receptor extracellular domain (ECD) relative to the plasma membrane (Figure 4C). A similar principle has been used previously to monitor epidermal growth factor receptor (EGFR) conformational shifts by FRET microscopy (Ziomkiewicz *et al*., 2013). Here, both ligands led to clear increases in TR-FRET signal across a wide concentration range. Efficacy was reduced with exendin-4-C16 (Figure 4C), in keeping with lower efficacy measurements for intracellular responses including recruitment of mini-G_s_ and β-arrestin-2 (see Figure 1), although an increase in potency was observed. The implications of these responses are not clear, as they could reflect differences in ECD movement, known to be a feature of GLP-1R activation as determined from structural and computational studies (J Zhang *et al*., 2019; Wu *et al*., 2020), but ligand-induced effects on membrane architecture cannot be ruled out.

To determine whether manipulation of membrane liquid order would differentially affect the signalling responses of exendin-4 *versus* exendin-4-C16, we used methyl-β-cyclodextrin (MβCD) to sequester cholesterol and increase membrane liquid disorder (Mahammad and Parmryd, 2015). MβCD treatment disproportionately reduced cAMP production by exendin-4-C16 compared to exendin-4 (Figure 4D). This might result from the lower G protein recruitment efficacy of exendin-4-C16 meaning that its ability to generate cAMP signals is more susceptible to uncoupling of G proteins from receptors as a result of disruption of membrane structure (Mystek *et al*., 2016). Similarly, exendin-4-C16-induced GLP-1R endocytosis, measured by TR-FRET, was more substantially affected by MβCD treatment than exendin-4 (Figure 4E).

### 3.5 Interaction of exendin-4-C16 with albumin and *in vivo* efficacy

The classical benefit of peptide acylation is to promote reversible binding to albumin in the circulation, thereby avoiding renal elimination. We previously showed that exendin-4-C16 and related analogues are >97% bound to proteins in human and mouse plasma (Lucey *et al*., 2020). In the present study we performed fluorescence correlation spectroscopy (FCS) to measure the interaction of both TMR-conjugated peptides with bovine serum albumin (BSA) (Figure 5A-C). All traces could be fitted to a 1×3D free diffusion model, indicating there is a single diffusing component. Diffusion coefficients for exendin-4-TMR and exendin- 4-TMR-C16, tested at a range of concentrations between 2.5 and 20 nM, were similar in the absence of BSA (Figure 5B). However, diffusion of exendin-4-TMR-C16 was markedly slowed in the presence of BSA, indicative of formation of peptide-albumin complexes. (Figure 5A, 5B). Exendin-4-TMR-C16 measurement reads were reanalysed by photon counting histogram (PCH) analysis with all reads fitting to a single-species PCH fit and the reported average molecular brightness unaffected by the addition of BSA (Figure 5C). These results support our interpretation of a slowing of diffusion coefficient as a result of exendin-4- TMR-C16 interaction with BSA rather than an accumulation of peptide, which would have resulted in an increase in average molecular brightness.

**Figure 5.**
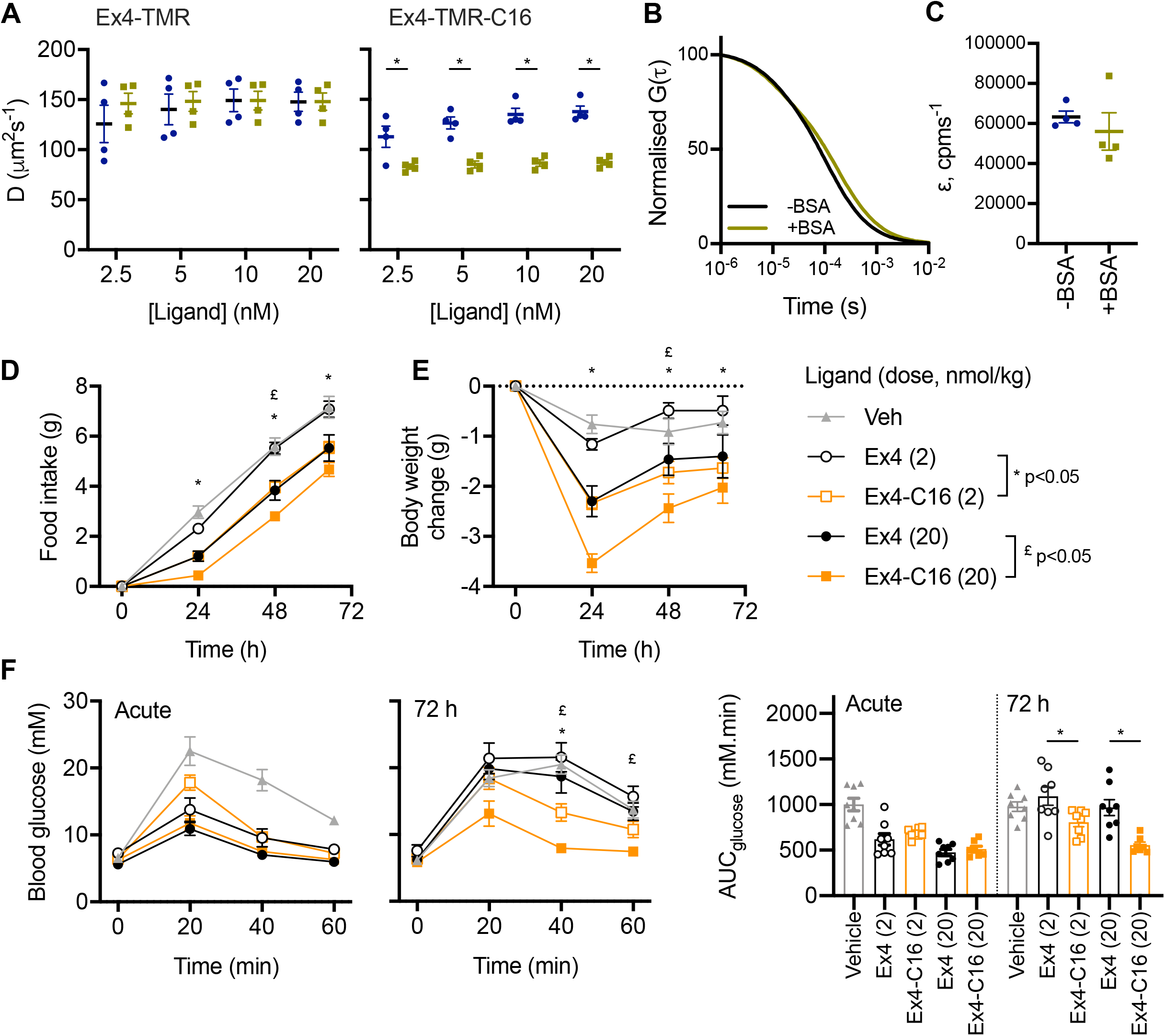
Albumin binding by fluorescence correlation spectroscopy and *in vivo* effects. (**A**) Diffusion coefficients, D (µm^2^s^-1^), of indicated concentration of indicated ligand with and without 0.1% BSA, with comparison by 2-way randomised block ANOVA with Sidak’s test. (**B**) Representative 1x 3D free diffusion fit for Exendin4-TMR-C16 (20 nM) autocorrelation curves with or without 0.1% BSA. The autocorrelation function, G(τ), is normalised to facilitate comparison of decay curves. (**C**) Molecular brightness, ε (counts per molecule per second), of EX4-TMR-C16 (20 nM) with and without 0.1% BSA. (**D**) Cumulative food intake in male HFD mice (n=8/group) after a single IP injection of the indicated agonist. Statistical comparisons between equimolar agonist doses by 2-way repeated measures ANOVA with Tukey’s test. (**C**) As for (B) but showing the effect on body weight. (**E**) IPGTT (2g/kg glucose) performed at the start and end of the 72-hour study. AUCs are compared by 1-way ANOVA with Sidak’s test for equimolar agonist doses. Data are shown as mean ± SEM with individual replicates shown in some cases. * p<0.05 by statistical test indicated.

Finally, the metabolic effects of exendin-4 and exendin-4-C16 were compared in mice in which glucose intolerance was first induced by high fat feeding for 2 months prior to the study. Exendin-4-C16 was previously shown to be detectable in plasma 72 hours after dosing (Lucey *et al*., 2020). Therefore, the effect of a single administration of each ligand, at two separate doses, was assessed. Both doses of exendin-4-C16 led to greater suppression of food intake and weight loss throughout the dosing period (Figure 5D, 5E). Moreover, both doses of the acylated ligand exerted a larger anti-hyperglycaemic effect in an intra-peritoneal glucose tolerance test performed 72 hours after dosing (Figure 5F).

## 4 Discussion

This study demonstrates how C-terminal acylation of exendin-4 affects several GLP-1R pharmacological properties that are relevant to its therapeutic effect. Whilst exendin-4-C16 showed minimal reduction in cAMP signalling potency, marked differences were observed for recruitment of key intracellular effectors and trafficking responses, which led to increased insulin secretion with prolonged incubations. We also evaluated the effect of peptide acylation on binding to membranes and albumin to more comprehensively describe the differences between these ligands. Overall, our study highlights the breadth of pharmacological parameters than can be affected by peptide acylation.

This work was prompted by our earlier observation that C-terminal acylation of biased GLP-1RAs based on exendin-4 results in reduced recruitment efficacy for β-arrestin and mini-G protein (Lucey *et al*., 2020). The present study confirms this observation, with a 60% reduction in maximum response recorded for β-arrestin-2 recruitment by nanoBiT complementation with exendin-4-C16. Whilst mini-G_s_ recruitment was also reduced, the impact on β-arrestin-2 recruitment was greater, with biased agonism confirmed by the operational model approach and on the basis of efficacy differences (Onaran *et al*., 2017). This adds to the growing body of evidence that G protein-favouring biased GLP-1RAs typically show reduced efficacy for recruitment of both G proteins and β tins (Lucey *et al*., 2020; Pickford *et al*., 2020; Fang, Chen, Pickford, *et al*., 2020). However, using a novel Nb37-based BRET approach to monitor activation of endogenous G proteins close to GLP-1R, we demonstrate here that exendin-4-C16 shows increases in G_s_ activation in spite of its lower mini-G_s_ recruitment efficacy. Whilst the dynamic range of this assay was low in our hands, making it challenging to apply to higher throughput screening efforts or to concentration responses, it could be adapted by the use of alternative fluorophores with greater signal separation from the nanoluciferase emission peak (Dale *et al*., 2019) or using complementation approaches (Inoue *et al*., 2019). Other approaches to monitor G protein activation have typically required overexpression of tagged G protein subunits (Masuho *et al*., 2015; Inoue *et al*., 2019; Zhao *et al*., 2020; Olsen *et al*., 2020), which may not totally replicate the physiological setting.

In line with other studies showing that lower efficacy biased GLP-1RAs tend to induce slower GLP-1R endocytosis (Jones *et al*., 2018; Fremaux *et al*., 2019; Lucey *et al*., 2020; Willard *et al*., 2020), we observed that GLP-1R internalisation was reduced with exendin-4-C16 compared to exendin-4. The automated microscopy approach we used to demonstrate this has certain advantages over lower throughput methods, by allowing responses to a wide range of ligands or, in this case, ligand concentrations to be monitored in parallel across several FOVs, characterising ligand effects in more detail and with greater statistical robustness. However, this method is unable to discriminate between GLP-1R clustering at the plasma membrane *versus* bona fide endocytosis events, although these are intrinsically linked, with the former occurring rapidly after ligand stimulation as a precursor to uptake into clathrin-coated vesicles (Buenaventura *et al*., 2019). The system could be adapted for use with alternative fluorescence approaches to monitor internalisation, e.g. using pH-sensitive SNAP-labelling fluorophores (Martineau *et al*., 2017), or potentially by adapting the optical system to provide surface-only illumination by total internal reflection (TIR) through waveguide photonic chips (Opstad *et al*., 2020).

We also applied a series of complementary proximity-based techniques based on both BRET and FRET to monitor GLP-1R redistribution between the plasma membrane and different endosomal compartments. Monitoring GLP-1R disappearance from the plasma membrane by DERET has been widely applied (Roed *et al*., 2014; Jones *et al*., 2018), and the use of the cleavable BG-Lumi4-Tb to monitor GLP-1R recycling in combination with a fluorescent antagonist ligand was recently described by our group (Pickford *et al*., 2020). We extended this approach here to detect GLP-1R translocation to late endosomes and lysosomes marked by the lysomotropic dye LysoTracker, facilitated by the spectral overlap of one of the Tb emission peaks with the excitation spectrum of LysoTracker DND99. In principle, a similar approach could be trialled with other fluorescent markers that accumulate in different subcellular compartments. Using a nanoluciferase tag at the GLP-1R C-terminus, we were also able to obtain BRET measurements of GLP-1R redistribution to early, late and recycling endosomes that corroborate the TR-FRET responses. This approach has been used recently to study the trafficking profiles of GLP-1R mono-agonists and dual GLP-1R/GIPR co-agonists (Fletcher *et al*., 2018; Novikoff *et al*., 2021), albeit using Renilla luciferase rather than the nanoluciferase we used in our study. The rank order of ligand-induced changes was consistent for “matched” TR-FRET and BRET approaches, with exendin-4-C16 showing reduced GLP-1R internalisation and lysosomal accumulation, but faster recycling. In our hands the TR-FRET approach resulted in clearer discrimination between ligand responses, which could reflect cell model differences, influence of the C-terminal NLuc tag for BRET, or assay/instrument sensitivity.

Importantly, we also observed differences in mini-G_s_ recruitment to plasma membrane *versus* early endosomes using the nanoBRET approach, providing some insight into compartmentalisation of GLP1-R signalling. Here, plasma membrane mini-G_s_ recruitment was somewhat reduced for exendin-4-C16 compared to exendin-4, which is compatible with results from our nanoBiT complementation assay demonstrating reduced global mini-G_s_ recruitment efficacy to GLP-1R for this ligand. However, recruitment of mini-G_s_ to Rab5-positive EEs was reduced to an even greater extent, which can be at least partly explained by the reduced rate of internalisation with this ligand, although differences in ability to maintain GLP-1R activation once internalised could also contribute. Indeed, the GLP-1R-Rab5 BRET signal was only marginally reduced with exendin-4-C16 *versus* exendin-4, whereas the mini-G_s_-Rab5 BRET response showed larger differences between ligands. However, differences in luciferase/fluorophore configuration between assays, as well as unknown effects of mini-G_s_ overexpression on GLP-1R pharmacology mean that further work will be needed to fully explain this observation. GLP-1R has been reported to generate signals from the endosomal compartment (Kuna *et al*., 2013; Roed *et al*., 2015; Girada *et al*., 2017; Fletcher *et al*., 2018), in line with many other GPCRs (Vilardaga *et al*., 2014), and this phenomenon is frequently claimed to be a mechanism for sustained cAMP signalling. However, while our results corroborate the existence of GLP-1R-associated endosomal signalling, they also suggest that sustained GLP-1R signalling, as indicated by cumulative insulin secretion over pharmacologically relevant timescales, is actually greater with the ligand (exendin-4-C16) with a reduced tendency to recruit mini-G_s_ to Rab5-positive endosomes. This raises questions about the relative therapeutic importance of maintaining an adequate pool of surface GLP-1Rs during prolonged stimulations *versus* aiming for maximal endosomal receptor activation. It remains to be investigated whether this phenomenon is exportable to other non-exendin-4-based GLP-1R agonist systems with differing tendencies to trigger GLP-1R lysosomal targeting. It should be emphasised that mini-G_s_ is a biosensor that detects G_s_-preferring GPCR conformations, rather than directly measuring signalling *per se*. Bystander BRET assays that monitor mini-G_s_ recruitment to different compartments therefore indicate the location of activated GPCRs, but additional approaches such as targeted cAMP or protein kinase A (PKA) FRET biosensors (Godbole *et al*., 2017; Fletcher *et al*., 2018) are needed to confirm whether this is associated with corresponding localised signalling responses.

The structural basis for the modified GLP-1R activation profile with exendin-4-C16 is not clear. The C-terminus of exendin-4, whilst not required for GLP-1R activation (Lee *et al*., 2018), plays an important role in GLP-1R binding (Doyle *et al*., 2003). Installation of a large acyl chain at the peptide C-terminus could potentially interfere with ligand binding. We observed a modest reduction in GLP-1R binding affinity, in keeping with this possibility. However, receptor activation efficacy was also reduced. Interestingly, the C-terminus of exendin-4 may be required to facilitate formation of high order GLP-1R oligomers through interaction with neighbouring GLP-1R protomers in *trans* (Koole *et al*., 2017). GLP-1R oligomerisation is reported to be required for full signalling responses (Harikumar *et al*., 2012). A further possibility is that the acyl chain could interact with the plasma membrane in a specific manner that interferes with GLP-1R activation. We found here that a fluorescently labelled exendin-4-C16 does indeed form interactions with model membranes whereas the equivalently labelled non-acylated exendin-4 does not. Corresponding measurements from living cells also suggested greater membrane interactions with the acylated ligand but did not support the possibility of localised activation of GLP-1R subpopulations situated in particular plasma membrane nanodomains.

As ligand-specific GLP-1R conformations are a further potential explanation for the ligand signalling efficacy differences, we devised a strategy to monitor movements between the receptor ECD and the plasma membrane, finding efficacy reductions with exendin-4-C16 that matched the reduced intracellular signalling responses also observed with this ligand. This approach may be more widely useful as a conformational sensor for other GPCRs, although it lacks the ability to detect changes at the receptor intracellular face that are needed to provide insights into conformational changes required for G protein interactions.

In summary, beyond the expected effects on binding to albumin, C-terminal acylation of exendin-4 led to changes in multiple pharmacological parameters relevant to downstream GLP-1R responses. These observations are more broadly relevant to drug discovery at peptide GPCRs for which ligand acylation is a valid approach to improve pharmacokinetics.

## Author contributions

Participated in research design: Lucey, Wang, Goulding, Minnion, Elani, Briddon, Bloom, Tomas, Jones.

Conducted experiments: Lucey, Ashik, Marzook, Wang, Goulding, Elani, Jones

Contributed new reagents or analytic tools: Marzook, Oishi, Broichhagen, Hodson, Jockers

Performed data analysis: Lucey, Ashik, Goulding, Elani, Jones

Wrote or contributed to the writing of the manuscript: all authors

## Funding acknowledgements

The Section of Endocrinology and Investigative Medicine is funded by grants from the Medical Research Council, Biotechnology and Biological Sciences Research Council National Institute of Health Research (NIHR) and is supported by the NIHR Biomedical Research Centre Funding Scheme. The views expressed are those of the authors and not necessarily those of the funders. This project was also supported by the Medical Research Council (MR/R010676/1, MR/N00275X/1 and MR/S025618/1, and MR/N020081/1), the European Federation for the Study of Diabetes, Diabetes UK (including grant 17/0005681), the Academy of Medical Sciences, Society for Endocrinology, The British Society for Neuroendocrinology, the Engineering and Physical Sciences Research Council, the European Research Council under the European Union’s Horizon 2020 research and innovation programme (Starting Grant 715884), UKRI Future Leaders Fellowship (MR/S031537/1), Fondation de la Recherche Médicale (Equipe FRM DEQ20130326503, EQU201903008055), Agence Nationale de la Recherche (ANR-19-CE16-0025-01, ANR-15-CE14-0025-02), Institut National de la Santé et de la Recherche Médicale (INSERM), and Centre National de la Recherche Scientifique (CNRS).

## Abbreviations

BRET: Bioluminescence resonance energy transfer
BSA: Bovine serum albumin
CyOFP: Cyan-excitable orange fluorescent protein
DERET: Diffusion-enhanced resonance energy transfer
FCS: Fluorescence correlation spectroscopy
FITC: Fluorescein isothiocyanate
GFP: Green fluorescent protein
GLP-1(R)(A): Glucagon-like peptide-1 (receptor) (agonist)
GUV: Giant unilameller vesicle
MβCD: Methyl-β-cyclodextrin
Nb37: Nanobody-37
POPC: 1-palmitoyl-2-oleoyl-sn-glycero-3-phosphocholine
T2D: Type 2 diabetes
TMR: Tetramethylrhodamine
TR-FRET: Time-resolved Förster resonance energy transfer

