## Supplementary Information for "Acylation of the incretin peptide exendin-4 directly impacts GLP-1 receptor signalling and trafficking"

**Contents:**

- **Supplementary Figure 1**
- **Supplementary Figure 2**
- **Supplementary Figure 3**
- **Supplementary Figure 4**

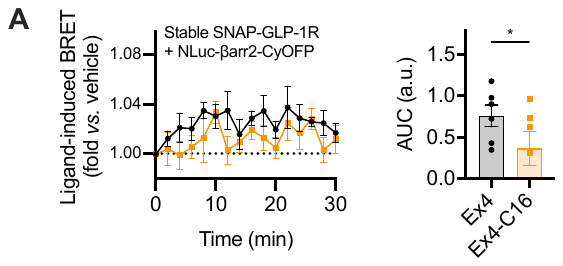

**Supplementary Figure 1. Pharmacological responses to exendin-4 and exendin-4-C16. (A)** Measurement of β-arrestin-2 activation in HEK293-SNAP-GLP-1R cells transiently transfected with NLuc-4myc-βarr2-CYOFP1 and stimulated with 100 nM agonist or vehicle, *n*=6, with AUCs compared by paired t-test. Data are shown as mean ± SEM with individual replicates shown for AUC graph. * p<0.05 by statistical test indicated.

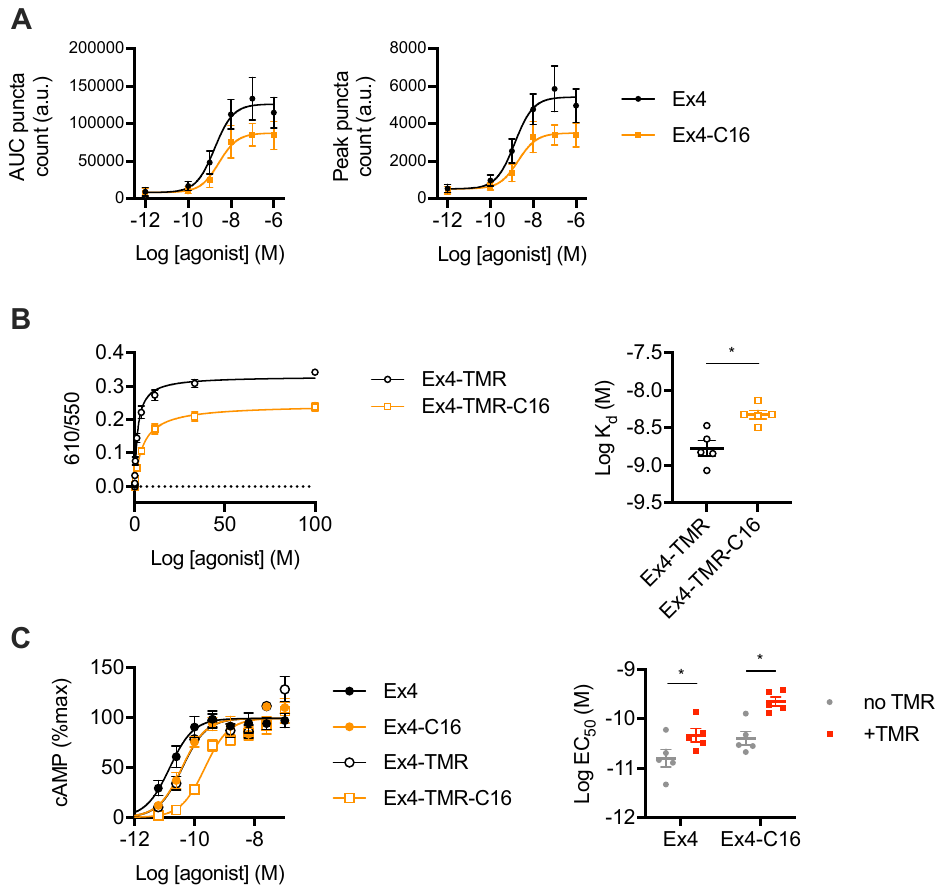

**Supplementary Figure 2. Trafficking responses to exendin-4, exendin-4-C16, and functional characterization of TMR-conjugates.** (**A**) Alternative quantification of endosomal puncta formation from Figure 2B, with concentration-dependent peak number of spots, or AUC across the entire 30-min stimulation period indicated with 3-parameter fit shown. (**B**) TR-FRET binding data for exendin-4-TMR and exendin-4-TMR-C16 in HEK293-SNAP-GLP-1R cells, *n*=5, with log K_d_ compared by paired t-test. (**C**) cAMP data for TMR-modified or not exendin-4 / exendin-4-C16, *n*=5, with 3-parameter fits shown. LogEC_50_ values are compared by 2-way randomised block ANOVA with Sidak’s test. Data are shown as mean ± SEM with individual replicates shown where possible. * p<0.05 by statistical test indicated

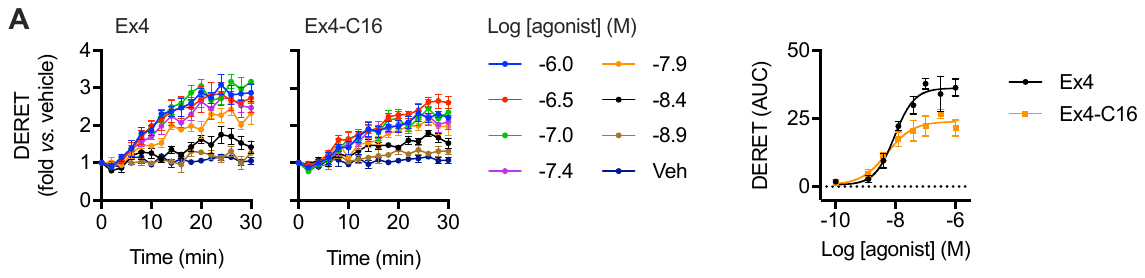

**Supplementary Figure 3. GLP-1R trafficking responses by DERET.** (**A**) GLP-1R endocytosis with exendin-4 and exendin-4-C16 measured by DERET in HEK293-SNAP-GLP-1R cells, *n*=5. Concentration response is quantified from AUC with 3-parameter fit shown. Data are shown as mean ± SEM.

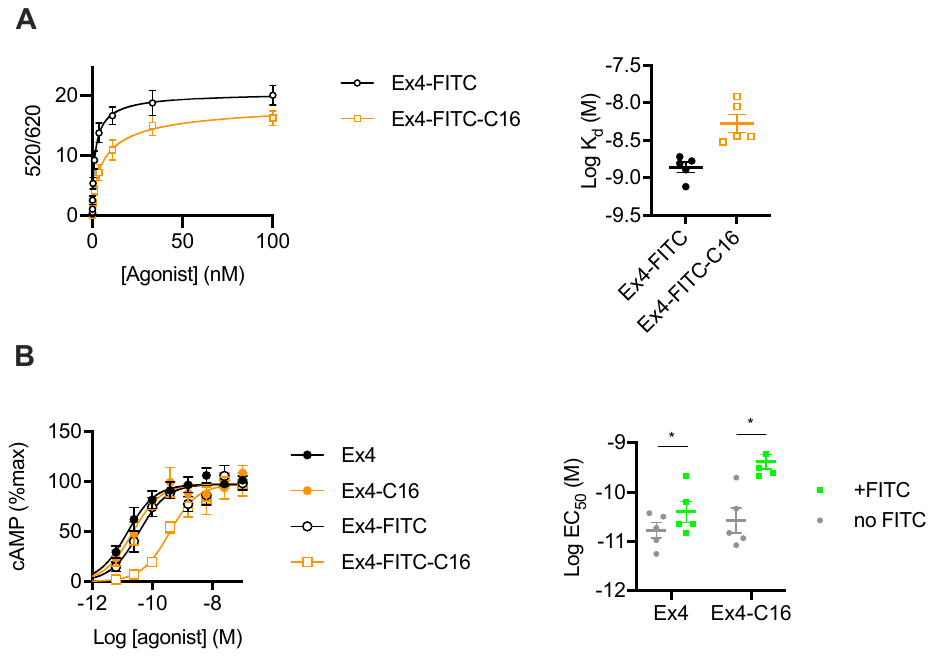

**Supplementary Figure 4. Exendin-4-FITC and exendin-4-FITC-C16 functional characterisation.** (**A**) TR-FRET binding data for exendin-4-FITC and exendin-4-FITC-C16 in HEK293-SNAP-GLP-1R cells, *n*=5, with log K_d_ compared by paired t-test. (**B**) cAMP data for FITC-modified or not exendin-4 / exendin-4-C16, *n*=5, with 3-parameter fits shown. LogEC_50_ values are compared by 2-way randomised block ANOVA with Sidak’s test. Data are shown as mean ± SEM with individual replicates where possible. * p<0.05 by statistical test indicated.
